# Comparative genomic analysis reveals a monophyletic cold adapted *Arthrobacter* cluster from polar and alpine regions

**DOI:** 10.1101/603225

**Authors:** Liang Shen, Yongqin Liu, Baiqing Xu, Ninglian Wang, Sten Anslan, Ping Ren, Fei Liu, Yuguang Zhou, Qing Liu

**Affiliations:** Key Laboratory of Tibetan Environment Changes and Land Surface Processes, Institute of Tibetan Plateau Research, Chinese Academy of Sciences, Beijing, 100085, China; CAS Center for Excellence in Tibetan Plateau Earth Sciences, Beijing, 100085, China; College of Urban and Environmental Science, Northwest University, Xian, 710069, China; Braunschweig University of Technology, Zoological Institute, Braunschweig, Germany; Institute of Microbiology, Chinese Academy of Sciences, Beijing, 100101, China; China General Microbiological Culture Collection Center (CGMCC), Institute of Microbiology, Chinese Academy of Sciences, Beijing 100101, China

**Keywords:** Cold adaptation, Genomic featrues, *Arthrobacter*, Polar and alpine regions

## Abstract

Decrease in the frequency of arginine and increase in lysine are the trends that have been identified in the genomes of cold adapted bacteria. However, some cold adapted taxa show only limited or no detectable changes in the frequencies of amino acid composition. Here, we examined *Arthrobacter* spp. genomes from a wide range of environments on whether the genomic adaptations can be conclusively identified across genomes of taxa from polar and alpine regions. Phylogenetic analysis with a concatenated alignment of 119 orthologous proteins revealed a monophyletic clustering of seven polar and alpine isolated strains. Significant changes in amino acid composition related to cold adaptation were exclusive to seven of the twenty-nine strains from polar and alpine regions. Analysis of significant indicator genes and cold shock genes also revealed that clear differences could only be detected in the same seven strains. These unique characteristics may result from a vast exchange of genome content at the node leading to the monophyletic cold adapted *Arthrobacter* cluster predicted by the birth-and-death model. We then experimentally validated that strains with significant changes in amino acid composition have a better capacity to grow at low temperature than the mesophilic strains.

**Importance:** Acquisition of novel traits through horizontal gene transfer at the early divergence of the monophyletic cluster may accelerate their adaptation to low temperature. Our study reached a clear relationship between adaptation to cold and genomic features and would advanced in understanding the ambiguous results produced by the previous studies on genomic adaption to cold temperature.

## Introduction

Separated by large distances and climatic barriers, high Arctic, Antarctic, and high alpine regions represent extreme cold environments, which have been successfully colonized by cold-adapted microorganisms (1-3). In the polar and alpine regions, cold-adapted microorganisms play a key role in biogeochemical transformations such as carbon, nitrogen, and iron, which have both local and global impacts (4-6); thus, it is important to understand the adaptation strategy of microorganisms in the extreme cold biome.

Temperature is a strong selective force that shapes the structure and function of microorganisms (7). Thriving in cold environments requires cold-adapted bacteria to synthesize enzymes that perform effectively at low temperatures, one of the major strategies is the modification to enzymes (8-10). Cold-adapted enzymes possess generally a number of amino acid changes that impart higher degree of structural flexibility (fewer salt bridges by reduce arginine and proline contents) and higher specific activity (more local mobility by increase asparagine, methionine and glycine contents) at low temperatures than their mesophilic counterparts (11). The increased low temperature activity of enzymes via change in amino acid composition was validated by analysis the structural, kinetic and microcalorimetric of cold adapted enzymes, e.g. aminopeptidase, β-lactamases, and dienelactone hydrolase (GaDlh) (12-14).

Comparative genomic analyses to examine differences in amino acid composition toward cold adaptation came through the study of two methanogenic Archaea, *Methanogenium frigidum* and *Methanococcoides burtonii* from Ace Lake, Antarctica (15). Proteins from these cold-adapted Archaea were characterized by a higher proportion of non-charged polar amino acids, such as glutamine and threonine, and a lower proportion of hydrophobic amino acids, particularly leucine (15). The amino acid shifts toward increased enzyme flexibility, which confers catalytic efficiency and contribute to cold adaptation can be identified in genomes of *Psychromonas ingrahamii* 37, *Exiguobacterium sibiricum* 255-15, *Psychrobacter arcticus* 273-4, *Shewanella* spp. and *Glaciecola* spp. (16-19). However, these changes were limited in *Colwellia psychrerythraea* 34H, *Planococcus halocryophilus* Or1, *Rhodococcus* sp. JG3, *Arthrobacter* spp., *Actinotalea* sp. KRMCY2, *Polaromonas* sp. Eur3 1.2.1, *Paenisporosarcina* sp. Eur1 9.01.10, *Methylobacterium* sp. AL-11 and *Kocuria* sp. KROCY2 (13, 20-24). Even, no changes in amino acid composition was identified in *Desulfotalea psychrophila*, *Psychroflexus torquis* and *Arcticibacter* spp. (25-27). Thus, these analyses, which attempt to find specific adaptations to cold environments by comparing the amino acid composition seem to have resulted in inconsistent outcomes (22, 28). Furthermore, Csp (cold shock protein) genes may vary widely across psychrophiles’ genomes, and are even absent in some strains, such as *Rhodoferax ferrireducens* T118 and *Methanolobus psychrophilus* R15. This indicates an ambiguous correlation between the number of Csp genes and cold tolerance (9, 29). It remains unknown why these genomic shifts toward cold adaptations are not common features that shared by strains isolated across the alpine and polar regions.

In the present study, we aimed to verify whether there are genomic features that could be conclusively identified by investigating the genomic patterns of *Arthrobacter* spp. that are isolated from permanently cold environments. These strains share close last common ancestry, and therefore differences observed between cold derived genomes and the reference genomes are more likely to be the result of cold adaptation (22, 30). Furthermore, we also investigated whether changes in the amino acid composition towards cold adaptation can promote growth at low temperatures.

## Materials and Methods

Sixteen *Arthrobacter* strains were isolated from the Tibetan Plateau (TP) using R2A medium at 4 °C, (Supplementary Table S1). Growth curves at various temperatures (25 °C, 5 °C and −1 °C) was measured in R2A broth. For the −1 °C temperature treatment, the temperature was maintained with ice–water mixtures and by controlling the ambient temperatures at 0 °C. The R2A broth remained liquid at −1 °C (27). Other temperature treatments were sustained using a constant-temperature incubator. To monitor growth, absorbance was measured at 600 nm on a Microplate Reader (MD SpectraMax M5). L. Stokes (31) have suggested a one week period of incubation at 0 °C in order to define how well microorganisms are adapted to cold. However, in the present study we performed 10 days of incubation at −1 °C. The growth curve tests performed on strains were deposited at the CGMCC (China General Microbiological Culture Collection Center) under the accession numbers *Arthrobacter* sp. N199823 = CGMCC1.16197, *Arthrobacter* sp. 4R501 = CGMCC1.16194, *Arthrobacter* sp. B1805 = CGMCC1.16193, *Arthrobacter* sp. 08Y14 = CGMCC1.16198 and *Arthrobacter* sp. 9E14 = CGMCC1.16188. Strains *Arthrobacter* sp. A3 (CGMCC1.8987), *A.* alpinus CGMCC1.8950, *A. globiformis* CGMCC1.1894 and *A. luteolus* CGMCC1.1218 were from CGMCC. The whole genome shotgun sequences were deposited at DDBJ/ENA/GenBank under the BioProject PRJNA421662.

The genomic DNA of the sixteen strains were extracted using TIANamp Bacteria DNA Kit (TIANGEN, Beijing) following to the manufacturer’s instructions. The concentration of genomic DNA was assessed with a NanoDrop spectrophotometer (2000c, Thermo Scientific, USA) and had an OD 260/280 ratio of 1.8–2.0. The DNA was stored in TE buffer (pH 8.0) for genome sequencing. Sequencing was performed using Illumina Hiseq 2000. Reads were assembled using SPAdes v3.11.1 with default options (32). As the algorithm is sensitive to sequencing errors, low-quality reads were filtered prior to *de novo* assembly using Fastp with default options (33).

Reference genomes were downloaded from NCBI in March 2017. The completeness of genomes was calculated using CheckM v1.0.7 with options lineage_wf, -t 16, -x fna (34). rRNA genes were called using RNAmmer (v.1.2) (35). Genomes with a completeness of less than 96% and lack of extractable full-length 16S rRNA reads were removed. The resulting set of 39 reference *Arthrobacter* genomes were used for further analysis along with the 16 genomes obtained in this study. Of the 55 genomes, 29 were from extreme cold polar and alpine regions, and 26 were from non-extreme-cold environments (Supplementary Table S1).

The air temperatures (2 m from surface) were downloaded from the European Centre for Medium-Range Weather Forecasts (ECMWF) ERA-Interim reanalysis Database (http://apps.ecmwf.int/datasets/data/interim-full-moda/levtype=sfc/) (36). Periods of 2008-2017 were selected from ERA-Interim with a spatial resolution 1.5°*1.5°. For the strains of *A. woluwensis* NBRC 107840, *A. woluwensis* DSM 10495 and *A. luteolus* NBRC 107841 isolated from human body we used 37 °C as the environment temperature.

All the genomes were annotated simultaneously in the present study with RAST (Rapid Annotation using Subsystem Technology) (37). Calculation of amino acid composition was carried out with the PERL script ‘aminoacidUsage.pl’ (38). One-way analysis of variance (ANOVA) was used to examine the differences in amino acids composition. Statistical significance was considered at α ≤ 0.05.

For gene family clustering, *Microbacterium* sp. (No.7) was chosen as the out-group. *Microbacterium* is one of the closest relatives to the *Arthrobacter* genus (39) and it is placed right at the lineage outside the *Arthrobacter*. In general, out-groups that closely related to the in-group species are better suited for phylogeny reconstruction than distantly related out-groups (40). Gene families were clustered using FastOrtho software (--pv_cutoff 1-e5 --pi_cutoff 50 --pmatch_cutoff 50) (http://enews.patricbrc.org/fastortho/). At the amino acid level using an E value of 10^−5^ and ≥ 50% global amino acid identity threshold, a total of 36, 699 orthologs were identified. Among these, 148 were universal to all the genomes sampled and 119 were single-copy orthologs. The number of single-copy identified in the present study is approximately consistent to the study of D. H. Parks et al. (41) which found 120 shared single-copy proteins in bacteria. The 119 mono-copy orthologs were then concatenated using custom-made PERL scripts. As a first step for a genome tree construction, the concatenated orthologous genes were aligned at the amino acid sequence level using Muscle software v3.8.31 (42). Non-conserved segments in the alignments were then trimmed using the Gblock v0.9b (43) to discard all gap-containing columns (-b1 = 50 -b4 = 5, other parameters were set as default). As a second step, probabilistic phylogenetic approaches were used to analyze the concatenation data (30, 366 sites) of the 119 orthologs. The PTHREADS version of RAxML v8.2.4 (-f a –x 12345 –p 12345 -#100 -m GTRGAMMAI) and IQ-TREE v1.6.0 (-b 1000 –m GTR+I+G4) were used to construct a Maximum Likelihood phylogenetic tree (44, 45). The MPI version of Mrbayes v3.2.6 (mcmc nchains = 16 burnin = 0.25, samplefreq = 100, Ngen = 10000000, lset nst = 6 rates = gamma) was used to construct a Bayesian phylogenetic tree (46). As the evolutionary models for different sites in multi-gene concatenated alignments may differ, PartitionFinder software v2.1.0 was used to determine the best-fit partitioning scheme for RAxML and Mrbayes (47) with default settings. The resultant trees were embellished with Adobe Illustrator CS6 and iTOL v3 (48).

Ordinations and statistical analyses were performed using the vegan package v2.4.4 (49) using R v3.3.3. Genes significantly associated with cold and temperate environments were calculated by Indicator Species Analysis as implemented in the R library labdsv (http://ecology.msu.montana.edu/labdsv/R/). Significance was calculated through random reassignment of groups with 1,000 permutations.

The 55 *Arthrobacter* genomes were used for ancestral reconstruction; the out-group species *Microbacterium* sp. (No.7) was not included because reconstruction of ancestral genome content using COUNT v9.1106 does not require out-group species (50). The COUNT software uses birth-and-death models to identify the rates of gene deletion, duplication, and loss in each branch and node of a phylogenetic tree. We used the pan-genome matrices (Supplementary Table S2) and the phylogenetic birth-and-death model implemented in COUNT, to reconstruct the ancestral genome content of *Arthrobacter* species. Ancestral history reconstruction was performed by posterior probabilities: one hundred rounds of rate optimization were computed with a convergence threshold of 10^−3^ prior to ancestral reconstruction, other parameters were set as default; Horizontally-transferred genes (HTgenes) were identified using a threshold of probability of gain higher value than 0.95 at the destination node and excluding gains occurring in the last common ancestor with a probability higher than 0.5, as suggested by Oliveira and colleagues (51).

## Results

### Distribution of Arthrobacter strains along their phylogenetic clade

We constructed phylogenetic clustering based on concatenated alignment of 119 orthologous to yield a high-resolution tree. The 55 *Arthrobacter* strains were clustered into three main lineages (lineage 1, 2 and 3 in Fig. 1a; Fig. S1). Strains isolated from polar and alpine environments were mixed in lineage 1 and 2 with the reference strains. For example, despite being isolated from different environments, strains *Arthrobacter* sp. Y81 from Tibetan Plateau (TP) lake, *Arthrobacter* sp. TB 26 from Antarctica marine sponge (52), *Arthrobacter* sp. FB24 from soil in Seymour, Indiana (53) and *Arthrobacter* sp. SPG23 from contaminated soil at the Ford Motor Company site in Genk, Belgium (54), clustered together in the lineage 1. Strains of *Arthrobacter* sp. Soil782 and *Arthrobacter* sp. H5 were located together in lineage 2, despite the former was isolated from plant material and the latter was isolated from Antarctic soil (24, 55). However, strains in lineage 3, in the middle of the phylogenetic tree (Fig. 1a), were all isolated from polar or alpine environments. These included strains of *Arthrobacter* sp. GMC3, *Arthrobacter* sp. A3, *Arthrobacter* sp. N199823, and *A. alpinus* ERGS4-06 (isolated from TP lake, permafrost, ice core and glacial stream water); *A. alpinus* R3-8 (isolated from Antarctic soil); *Arthrobacter* sp. PAMC 25486 (isolated from Arctic soil) and *A. alpinus* DSM22274 (isolated from alpine soil).

**Figure 1.**
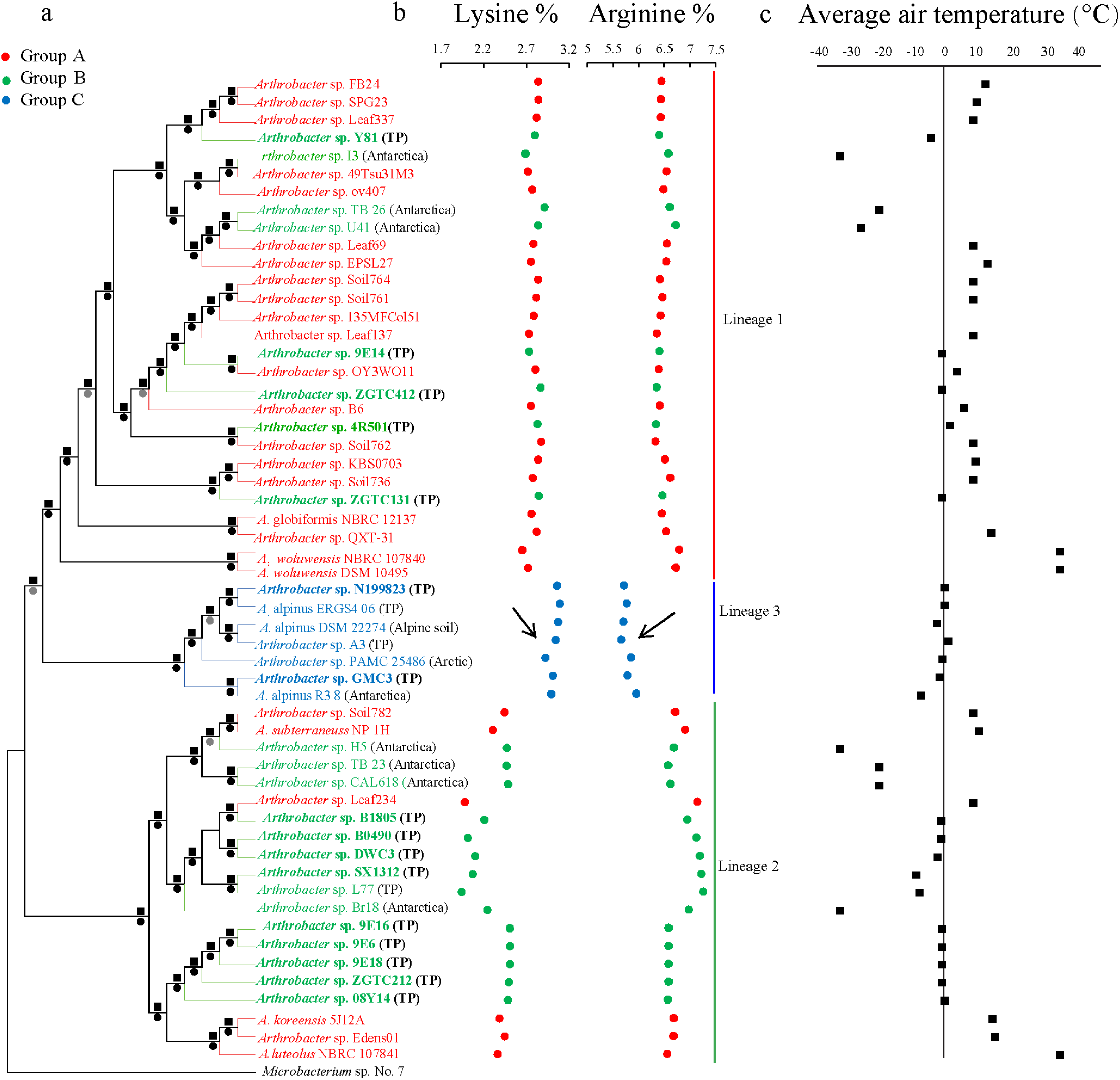
(a) Phylogenetic clustering of *Arthrobacter* strains based on concatenated alignment of 119 orthologous proteins using RAxML, numbers at nodes indicate posterior probability/bootstrap percentages by MrBayes and RAxML. (b) Relative composition (frequency compared to the total amino acids present) of lysine and arginine in protein sequences of *Arthrobacter* proteomes, arrows highlighted significant change in the relative composition of arginine and lysine. (c) Average annual air temperature of the strains (from 2008 to 2017), the environment temperature of the polar and alpine strains was significantly lower than that of the references strains (*P* < 0.005, F = 42.1).

Based on the clustering results, we classified these 55 strains into three groups for comparative genomic analysis: group A comprised the 26 reference strains isolated from non-extreme-cold environments (e.g. rhizosphere soil, plant leaf surface and blood culture out of polar or alpine) in lineages 1 and 2; group B comprised 22 strains isolated from cold environments (polar or alpine) in lineages 1 and 2; group C comprised 7 strains isolated from cold environments in lineage 3 (Fig. 1a). We detected a decrease in the frequency of arginine and an increase in lysine, which occurred exclusively to the genomes of strains belonging to group C (Fig. 1b). The air temperatures (2 m from surface) of the polar and alpine strains habitats was significantly lower compared with the references strains habitat (ANOVA, *P* < 0.005, F = 4.052, Fig. 1c).

### Strain growth at different temperatures

Strains isolated from Tibetan Plateau showed variable growth patterns, but in general grew better at low temperatures (5 °C, −1 °C, Fig. 2) than the references strains. Strains in group A and B grew faster than strains in group C in exponential phase before 36 h at 25 ℃ (Fig. 2a). At 5 °C, strains in group C tended to grow faster than strains in group A and group B in exponential phase before 144 h except strain *Arthrobacter* sp. 4R501; strains in group B grew faster than strains in group A (Fig. 2b). At −1 °C, strains in group C tended to grow faster than all the strains in group A (growth was inhibited) and group B in exponential phase before 240 h (mean OD600 of group A, B and C was 0.00, 0.02 and 0.25 at 240 h, respectively, Fig. 2c). The result is in consistent with the study on snow-bacteria of the Tibetan Plateau which revealed the adaption to cold environments was the result from the expansion of their minimum growth-temperature (56).

**Figure 2.**
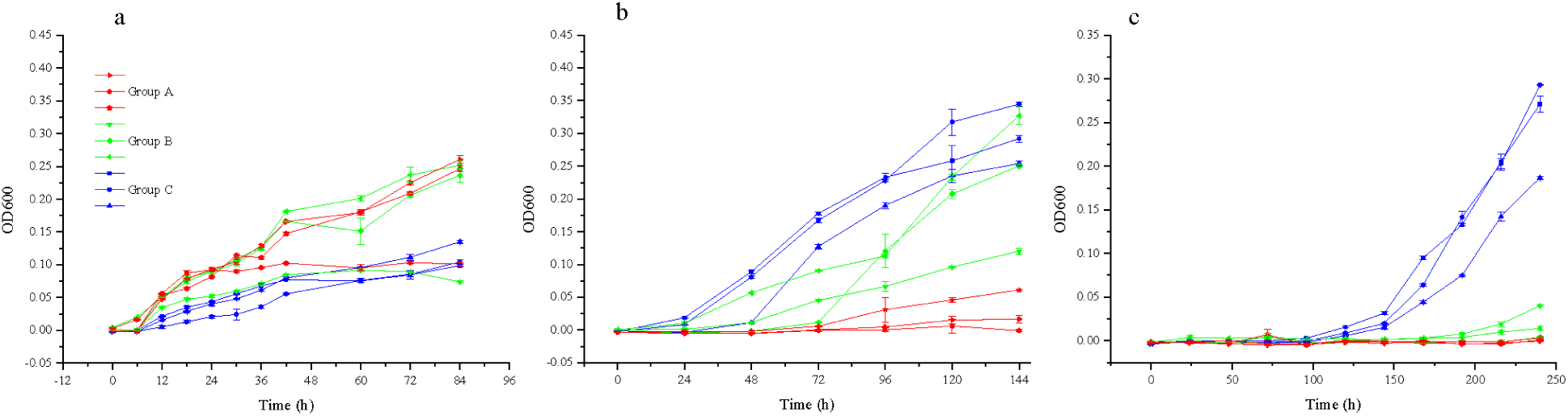
Growth curves for the Group A (*A. luteolus*, *A. globiformis* and *A. subterraneus*), group B (*Arthrobacter* sp. 4R501, *Arthrobacter* sp. 9E14 and *Arthrobacter* sp. 08Y14), group C (*A. alpinus, Arthrobacter* sp. A3 and *Arthrobacter* sp. N199823) grow at (a) 25 °C, (b) 5 °C and (c) −1 °C.

### Features in amino acid composition

The pattern of amino acid distribution in *Arthrobacter* spp. displayed an overall similar trends in their genomes, with alanine being most abundant, followed by leucine, glycine, and valine, while methionine, tryptophan and to a lesser extent cysteine were infrequent. However, compared against reference genomes, we found a decrease in the frequency of arginine and an increase in lysine have occurred exclusively in the genomes of strains belonging to group C. The shift in composition of arginine and lysine is closely related with survival strategies of psychrophiles (11). Then, we calculated the composition of the twenty common amino acids in whole proteins to determine the differences in their frequencies between the three groups (A, B and C). For group C, significant changes in twenty amino acids were apparent (ANOVA, *P* < 0.005, F = 4.1491, Fig. 3a). Of the amino acids that increased in group C, one is positively charged (lysine) and seven are uncharged (tryptophan, threonine, serine, asparagine, methionine, isoleucine and phenylalanine) (Fig. 3a). Of the amino acids that decreased in group C, three are charged (arginine, glutamic acid and aspartic acid), and three are uncharged (proline, glycine and alanine) (Fig. 3a). By exhibiting a number of amino acid changes that impart higher degree of structural flexibility (fewer salt bridges by reduce arginine, glutamic acid, aspartic acid and proline contents) and higher specific activity (more local mobility by increase asparagine, methionine and lysine contents) (11, 57), the enzyme activities of group C strains were predicted to be increased. The resulted increase in flexibility and decrease in thermodynamic stability were consistent with the experimental data that strains in group C grew better at low temperatures while weakly at a higher temperature compared to the reference strains. For group B, there were only changes in glycine (ANOVA, decrease with *P* = 0.0245, F = 4.052) and threonine (increase with *P* = 0.0366, F = 4.052) (Fig. 3b). The differences in arginine, lysine and proline composition were not significant in group B.

**Figure 3.**
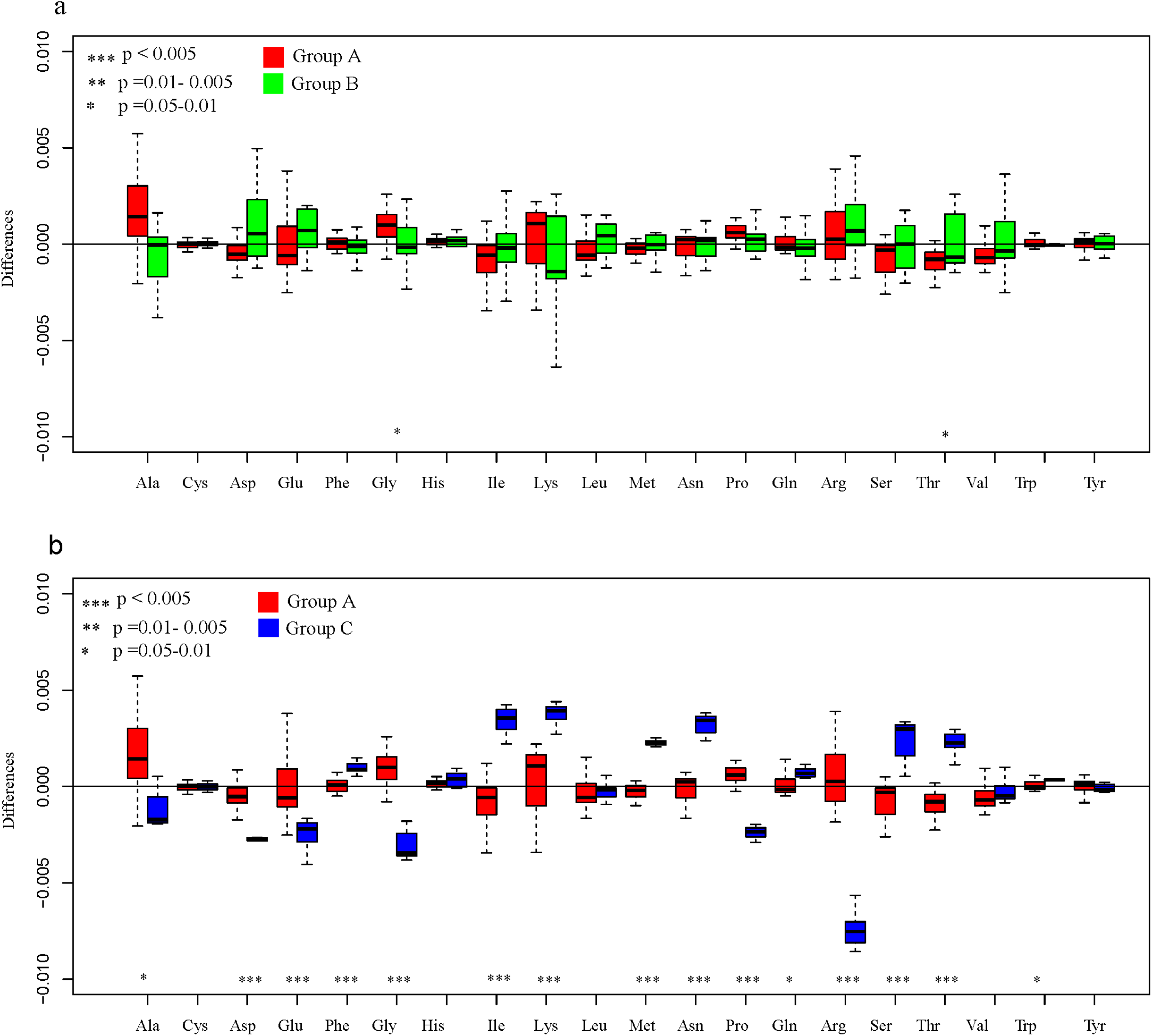
Changes in overall amino acid frequencies between (a) group A and B, (b) group A and C.

### Differences in the distribution of gene families between Arthrobacter strains

We performed a multivariate assessment of gene composition classified at the level of function. Ordination of functional genes using two-dimensional nonmetric multidimensional scaling (NMDS) revealed a clear separation of group C, while group A and B were not clearly separated (Fig. 4a). The PERMANOVA analysis with 1000 permutations (58) showed significant difference between strains of group A and C at a functional level (Fig. 4a, *P* = 0.001, F = 6.492). To remove the potential difference introduced by the distance between lineage 1 and lineage 2, we performed the PERMANOVA analysis between group A and B separately in lineage 1 and lineage 2. The results revealed no significant differences between cold-environment derived strains and the reference strains in lineage 1 and lineage 2 (*P* = 0.505 and *P* = 0.171, respectively).

**Figure 4.**
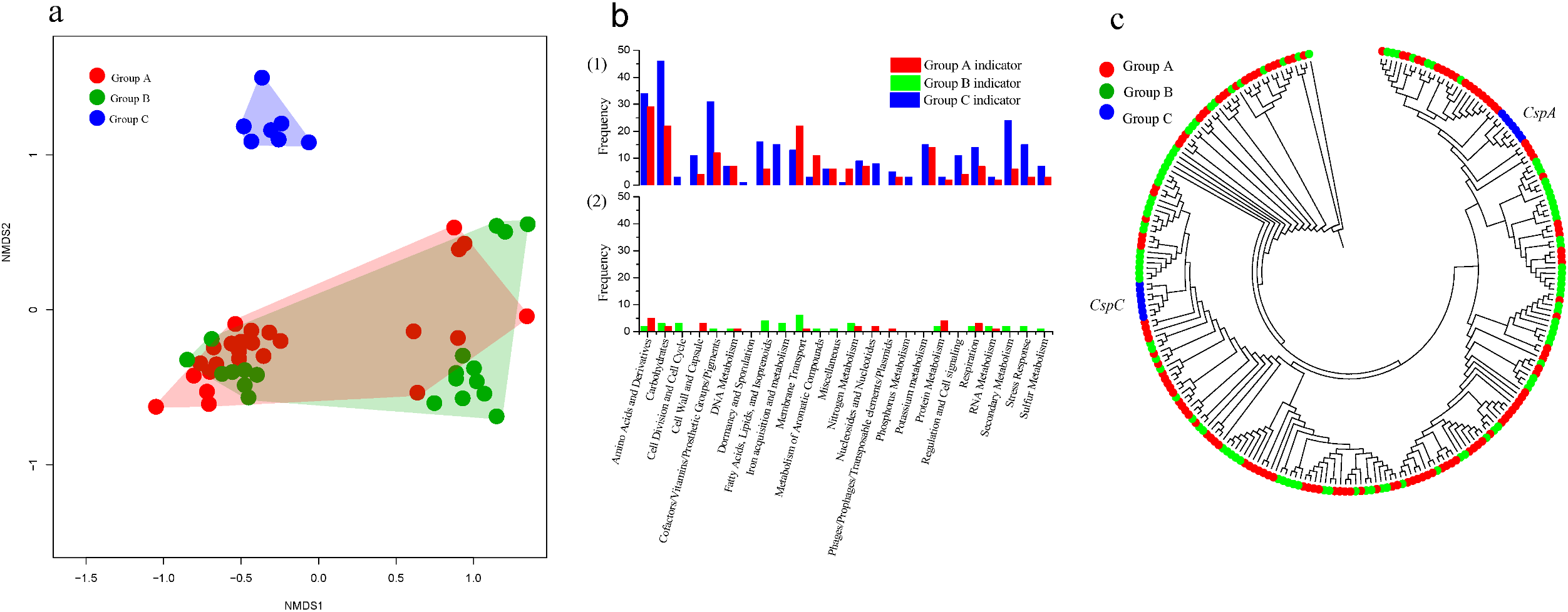
(a) Ordination of functional genes (classified at the level of function) using two-dimensional nonmetric multidimensional scaling (NMDS). Shading shows clear separation of group C (PERMANOVA, *P* = 0.001). (b) The frequency of significant indicator genes (*P* < 0.05) in (1) a comparison of group A and C strains, (2) a comparison of group A and B strains. (c) Phylogenetic relationships of genes related to cold shock; a close clustering between *cspA* and *cspC* from genomes of group C shows that these genes are monophyletic and can be clearly separated from those from group A and B which have an interleaved clustering of genes.

We further used indicator gene analysis to statistically define the characteristic genes contributing to the differences between the three groups. When group A was compared with group C, a total of 176 and 304 significant functional indicators were found in group A and C, respectively (Fig. 4b, ANOVA, *P* < 0.05, supplementary Table S3). In contrast, when group A was compared with group B, only 25 and 40 significant functional indicators were found, respectively (Fig. 4b, supplementary Table S3). The group C indicator genes were mainly affiliated to the functional category of Carbohydrates, Amino Acids and Derivatives and Cofactors/Vitamins/Prosthetic Groups/Pigments (Fig. 4b, supplementary Table S4).

### Cold shock genes of Arthrobacter strains

All of the genomes in group C contained two predicted cold shock genes (one copy of *cspA* and one *cspC*). All of the strains in group A and B had an extra 1 to 5 copies of *cspA*, except strain *Arthrobacter* sp. H5 (Supplementary Table S2). We generated alignments and a maximum likelihood tree for cold shock genes. The phylogenetic analyses showed that *cspA* and *cspC* genes of strains in group C were monophyletic, while cold shock proteins (Csps) from group B were polyphyletic, interleaving with group A (Fig. 4c).

### Ancestral reconstruction of Arthrobacter strains and the dynamics of genome content

We obtained a phylogenomic tree of the 55 *Arthrobacter* strains to reconstruct the ancestral genome content using three different phylogenomic approaches (IQ-TREE, RAxML and MrBayes; Fig. 5a and Fig. S1). The phylogenetic birth-and-death model imposed on the phylogenomic tree revealed a steady trend towards genome expansion since the most recent common ancestor indicated by N54 (~ 2,460 gene families, Fig. 5a). The extant *Arthrobacter* genomes (2,867 to 4,521 gene families, average ~3,500 gene families, Fig. 5a) exhibited a complicated evolutionary path to net genome expansion. Our attention was mainly focused on nodes of N27, N33 and N53 leading to lineage 1 and lineage 2 and lineage 3 (refer to group C), because strains in group C could be distinguished from group A with respect to overall amino acid composition, NMDS analysis and indicator genes. In contrast, these differences could not be detected between group A and B. Our results showed that node N33 that gained 876 genes, was more divergent than nodes N27 and N53 which gained 131 and 91 genes, respectively (Fig. 5b). The average genes gained per branch, showed the same trend (N33 = 125, N27 = 5 and N53 = 4), indicating that the significant difference between N33 and others was not due to the lineage size. The number of genes lost at node N33 was also higher compared to N27 and N53 (the number of lost genes: 362, 47 and 40, respectively; Fig. 5b). Almost half (429 of 876) of these genes were identified as HTgenes for node N33, while none of them were identified as HTgenes at N27, N33 and N53. Genes gained at node N33 were related to cofactors/vitamins/prosthetic groups/pigments (5.6%), membrane transport (3.7%), carbohydrates and amino acids (3.7%) and derivatives (3.7%), but the function of large proportion of gene remained unknown (65.2%) (Fig. 5c).

**Figure 5.**
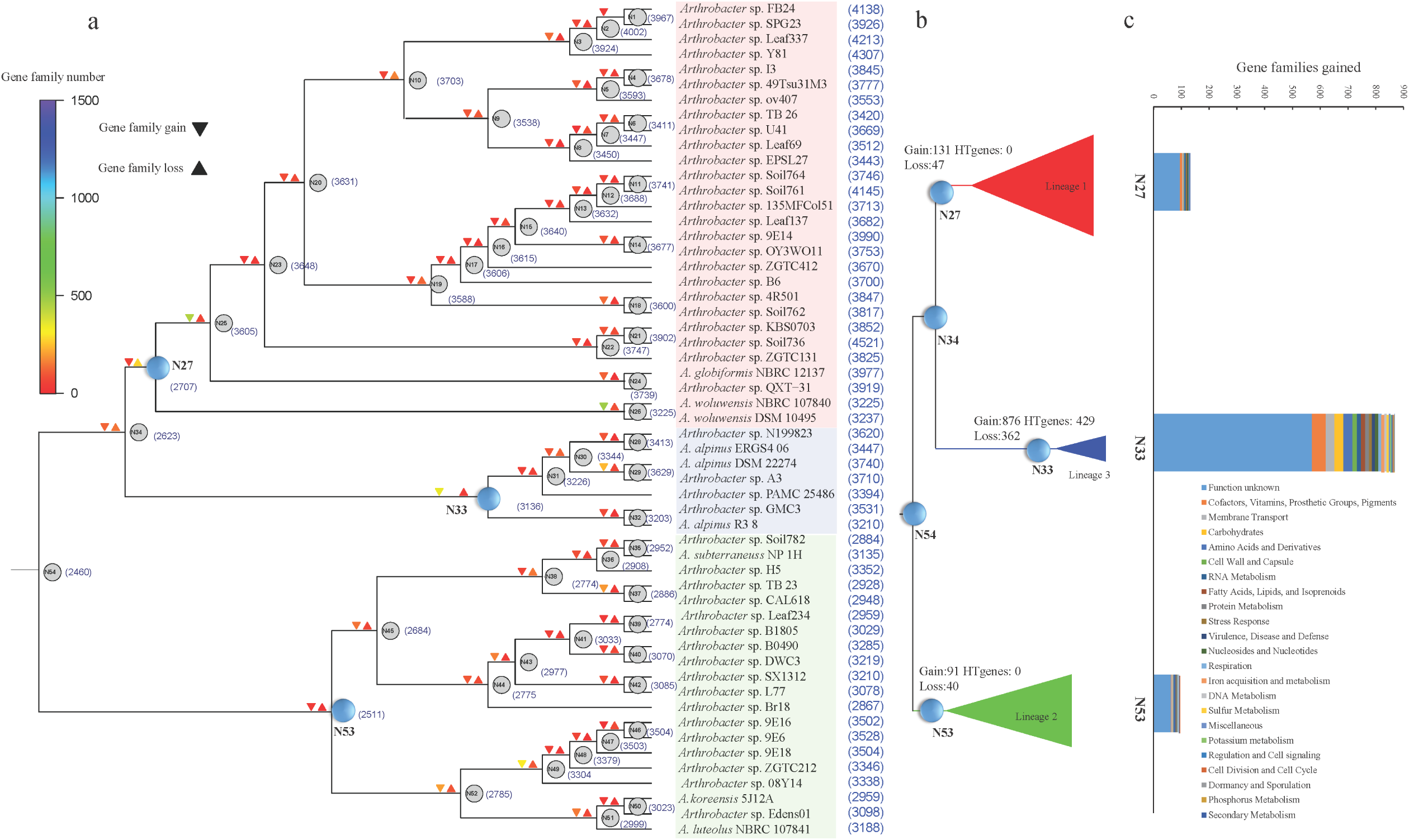
(a) Ancestral genome content reconstruction using COUNT software. The reconstruction is based on the RAxML *Arthrobacter* tree. The log-scale color coding represents the number of reconstructed gain and loss events for each lineage. Numbers in parentheses are predicted gene numbers for ancestral nodes and observed gene numbers for extant lineages. (b) Condensed and linearized maximum likelihood cladogram, showing the genome dynamics on the branches leading to nodes N27, N33 and N53. (c) Bar plot showing the distribution of genes families gained at N27, N33 and N53.

## Discussion

In the current study, we found that not all *Arthrobacter* strains isolated from polar and alpine regions exhibited significant detectable changes in the genome composition. Although changes in amino acid composition towards cold adaptation can be widely identified in psychrophilic strains (9, 11), strains in our group B which were isolated from cold environments and able to grow at 5 °C, exhibited no significance in changes in amino acid composition. This has also been the case of Antarctic *Arthrobacter* isolates, for example, strains *Arthrobacter* sp. FB24, *Arthrobacter* sp. Br18 and *Arthrobacter* sp. H5 were not located in lineage 3 in Fig. 1a, in which no remarkable genomic features were identified (24). Thus, in certain cases, amino acid shifts are limited and even no changes in amino acid composition can be identified, despite the strains were able to grow at low temperatures. However, the trends in amino acid composition in group C exhibited adaptations to cold environments in terms of genome-wide amino acid composition, which were consistent with those of cold adapted bacteria (9, 11). Thus, conserved genomic traits were exclusive to a certain group in *Arthrobacter*.

Strains in group C shared more genomic traits with the established rules for protein adaptation to cold than that of in group B (15, 17, 18, 57, 59). Thus, strains in group C may represent a “better adapted” genome type which were better adapted to their native cold habitats, this is supported by the faster growth of strains in group C at 5 and –1 °C than strains in group A and B. The wide abundance of group C indicator genes in the functional categories revealed that psychrophilic lifestyle is most likely conferred by a collection of synergistic changes in overall genome content rather than a unique set of genes (23). Grouping of bacteria isolated from cold environments based on their growth pattern at low temperature and phylogenetic clustering helped in the identification of conserved genomic traits of cold adaptation.

Cold shock proteins (Csp) regulate the cold shock response and play a critical role in bacterial growth at low temperatures (Jones and Inouye, 1994; Hébraud and Potier, 1999). Psychrophiles vary widely in the number of Csp genes present in their genomes, indicating a weak correlation between the number of Csp genes and cold tolerance (9). In the present study, we did not find any increase in the number of Csp genes in group C, which contains strains that grew well at −1 °C. On the contrary, strains in group C had fewer Csp genes than those in groups A and B despite the group C strains exhibiting faster growth rates atlow temperatures. Thus, increase in number of Csp genes may not be the strategy for cold adaptation of *Arthrobacter* spp. Strains in group C shared conserved genomic traits to cold adaptation and the clustering of these strains in one monophyletic lineage suggests that they all have evolved in cold environments and possess similar strategies to remain active and survive low/freezing temperatures. The dynamic historical pattern of *Arthrobacter* genomes is concordant with the emerging view that genomes evolve through a dynamic process of expansion and streamlining (60-62). However, the evolution of a cluster of strains which can be clearly separated from their relatives has rarely been studied. Our results showed that the N33 node (which leads to the strains in group C) exhibited early vast genome dynamics, which may play an important role in promoting the growth of these strains at low temperatures. The result is consistent with the study of Allen et al. (2009), which revealed the genome plasticity of *M. burtonii* that have enabled adaptations to cold environments. Also, an inter-order horizontal gene transfer event enabled the catabolism of compatible solutes by *Colwellia psychrerythraea* 34H, which provided a selective advantage in cold (63). This dynamic genome pattern is also in agreement with the general pattern of virtual, higher taxa in Archaea and *Streptococcus* genomes, which have the key mechanisms to help related taxa inhabit new niches (60, 61, 64). Because generation time in cold environments is generally longer, horizontal gene transfer may be more effective for acquiring beneficial traits rapidly (26, 65-68). This is experimentally validated by our results that species belonging to group C, which exclusivity exhibited vast dynamic in genome content, have a better capacity to grow at low temperature than their cold environment isolated counterparts.

Based on the result of reconstruction of most recent common ancestor, it is suggested that group C strains may be cold evolutionary legacies. Cold adaptations are superimposed on pre-existing microorganisms and the temperature-dependent distribution of bacteria may not result from widespread contemporary dispersal but is an ancient evolutionary legacy, as revealed by evolutional analysis of cold desert cyanobacteria and thermal traits of *Streptomyces* sister-taxa (69, 70). Strain *A. alpinus* DSM 22274 in group C is a new species isolated from Alps (71). Many other new bacteria species have been described from polar and alpine regions further suggesting that the level of cold origin taxa could be considerable, and these may represent endemic species (72-79).

## Conclusions

Changes in genome composition and obtaining new genes via horizontal gene transfer may not essential for bacteria to survive in cold environments. However, for strains belonging to group C in our study, their adaptation to cold is accelerated by the acquisition of novel traits through horizontal gene transfer (51). Our results indicate that growth at 5 °C may not require significant changes in genome content, but genomic modification seems to be essential for *Arthrobacter* spp. to grow well at subzero temperature (−1 °C in this study). We found that significantly conserved genomic traits could be detectable across the cold adapted strains that growth quickly at subzero temperature.

## Acknowledgements

This study was financially supported by the National Natural Science Foundation of China (Grant Nos. 41701085 and 41425004). Dr. Wei Zhu from the Beijing Institute of Genomics CAS is thanked for his bioinformatics assistance. Dr. Qilong Qin from Shandong University is thanked for his valuable suggestions.

